## Supplementary material for "Suspicions of two bridgehead invasions of *Xylella fastidiosa* subsp. *multiplex* in France": Sup table: Description of additionel supplementary files.docx

**Description of Additional Supplementary Files**

**File name: Supplementary Data 1**

**Description:** List of the 82 X. fastidiosa genome sequences used in this study.

**File name: Supplementary Data 2**

**Description:** "List of the X. fastidiosa strains used in this study and their VNTR allelic profile.

* CFBP refers to strains from the CIRM-CFBP (https://www6.inra.fr/cirm_eng/CFBP -Plant-Associated-Bacteria); LSV to strains isolated by the Plant Health Laboratory at ANSES, † NA: not available, ‡ the 16 strains used as “panel test” to set up the MLVA scheme, ¥ MLVA performed in extracted DNA instead of boiled suspension."

**File name: Supplementary Data 3**

**Description:** List of the 396 X. fastidiosa subsp. multiplex-infected French plant samples used in this study and their VNTR allelic profile. * 13 pairs of isolated strains and the DNA extracted from the same original plant sample, † the 16 samples for which DAPC clustering differed over the 20 runs, ‡ the samples collected by the DGAL in the framework of the national official surveillance strategy.

**File name: Supplementary Data 4**

**Description:** Summary of private allele frequencies of ST6 and ST7 samples.

**File name: Supplementary Data 5**

**Description:** Genetic differentiation of the clusters of X. fastidiosa subsp. multiplex estimated by RST (bellow the diagonal) and FST (above the diagonal) pairwise comparisons for A) the DAPC k=4 groups; B) the three ST6 clusters used in ABC analyses; C) the three ST7 clusters used in ABC analyses. All pairwise population comparisons were significantly different (P<0.05) after 1,000 permutations.

**File name: Supplementary Data 6**

**Description:** Hierarchical AMOVA for A) ST6 X. fastidiosa subsp. multiplex DiyABC groups and B) ST7 X. fastidiosa subsp. multiplex DiyABC groups.

**File name: Supplementary Data 7**

**Description:** Percentage of votes obtained for each scenario using abcrf. A) results of the ST6 bottom-up analyses; B) results of the ST6 top-down analyses; C) results of the ST7 bottom-up analyses; D) results of the ST7 top-down analyses.

**File name: Supplementary Data 8**

**Description:** Distribution of prior parameters used for the DiyABC analyses.
