## Supplementary material for "Suspicions of two bridgehead invasions of *Xylella fastidiosa* subsp. *multiplex* in France": Sup mat and meth

**This PDF file includes:**

Supplementary Text for materials and method

Figs. S1 to S12

Tables S1 to S9

### 1. Supplementary material and methods

#### 1.1 DNA extraction and whole-genome sequencing

Fifty-five strains were sequenced in this study using Illumina HiSeq X 4000 or PacBio technology (Supplementary data 1). For the 48 strains sequenced by Illumina HiSeq X, highly concentrated bacterial suspensions were prepared from fresh cultures in 5 mL of ultrapure sterile water. The genomic DNA was extracted using the Wizard® Genomic DNA Purification Kit (Promega), following the manufacturer's recommendations. DNA was rehydrated in 100 µL of ultrapure sterile water. Quality and quantity of extracted DNA were verified using a NanoDrop ND-100 Spectrophotometer (Thermofisher) and a Qubit Fluorometer 1.0 (Invitrogen). Library preparation and sequencing were performed on Illumina HiSeq 4000 platform (BGI, Hong Kong). For the seven strains sequenced using the PacBio technology, DNA was extracted from fresh cultures in 10 mL of ultrapure sterile water following Mayjonade *et al.*'s recommendations[1]. Library preparation and sequencing were performed at the GeT-PlaGe core facility, INRAE Toulouse, according to the manufacturer's instructions "Shared protocol-20kb Template Preparation Using BluePippin Size Selection system (15kb size Cutoff)". At each step, DNA was quantified using the Qubit dsDNA HS Assay Kit (Life Technologies). DNA purity was tested using the NanoDrop (Thermofisher) and size distribution and degradation assessed using the Fragment analyzer (AATI) High Sensitivity Large Fragment 50kb Analysis Kit. Purification steps were performed using 0.45X AMPure PB beads (PacBio). 10 µg of each DNA was purified then sheared at 40 kb using the megaruptor1 system (diagenode). Using SMRTBell template Prep Kit 1.0 (PacBio), a DNA and END damage repair step was performed on 5 µg of each sample. Then blunt hairpin adapters were ligated to the libraries. The libraries were treated with an exonuclease cocktail to digest unligated DNA fragments. A size selection step using a 10kb cutoff was performed on the BluePippin Size Selection system (Sage Science) with 0.75% agarose cassettes, Marker S1 high Pass 15-20 kb. Conditioned Sequencing Primer V2 was annealed to the size-selected SMRTbells. The annealed libraries were then bound to the P6-C4 polymerase using a ratio of polymerase to SMRTbell at 10:1. Then after a magnetic bead-loading step (OCPW), 7 SMRTbell libraries were sequenced on 7 SMRTcell on RSII instrument at 0.2 nM with a 360 min movie.

#### 1.2 MLVA

##### 1.2.1 Design and development of the VNTR-13 scheme

The complete genome sequence of the M12 strain of *X. fastidiosa* subsp. *multiplex* isolated from *Prunus dulcis* in the USA in 2003 (GenBank accession number NC_010513.1) was screened for the presence of candidate TR loci using Tandem Repeat Finder 4.09 (https://tandem.bu.edu/trf/trf.html). Parameters were set as follows: total length in a range of 50–500 bp and TR length ≤ 44 bp. Other parameters were set as default. The identified loci were analyzed for (i) presence in 10 *X. fastidiosa* subsp. *multiplex*[2] using BLASTn, (ii) absence of redundancy with the previously published set of VNTR loci[3,4], and (iii) existence of polymorphism in the 10 *X. fastidiosa* subsp *multiplex* genome sequences by manual checking. Three novel VNTR loci (XFSSR-37, XFSSR-40 and XFSSR-58) were retained for further analyses to complete the set of 10 previously published VNTR loci (nine VNTR loci previously designed by Lin *et al*. (2005) and one VNTR locus designed by Francisco *et al*. (2017), Table 1). The 150 bp flanking regions of these VNTR loci were extracted from the 10 genome sequences of *X. fastidiosa* subsp *multiplex*. Oligonucleotide primers were designed in their homologous segments using Primer3 2.3.4 [5]. The specificity of all 13 VNTR primer pairs was tested *in silico* using PrimerSearch (Val Curwen, Human Genome Mapping Project, Cambridge, UK) on the 154,478 bacterial Whole Genome Shotgun (WGS) sequences available in the NCBI database (as on August 22, 2018). The primer pair sets were tested using Amplify to verify the absence of dimer and cross-amplification with other bacterium species[6].

##### 1.2.2 Optimization on artificially infected plant of the VNTR-13 scheme

Primers were tested by PCR using 16 strains of *X. fastidiosa* as a “panel test” (Supplementary data 2). Primer pair multiplexing was done according to the annealing temperature. Finally, all forward primers were labeled with one of the fluorescent dyes (6-FAM, HEX, ATTO550 and ATTO565) (Eurofins) at their 5’ end. To optimize VNTR amplification protocol for direct use in plant material, the efficiency of two Taq polymerases (Platinum™ Taq DNA Polymerase, Invitrogen, and GoTaq® G2 Flexi DNA Polymerase, Promega) were compared, as well as the impact of the DNA volume (1 µL and 2 µL) added to the reaction mix on ASSR-9, ASSR-16 and GSSR-4 VNTR loci amplification. Tests were run on DNA extracted from maceration of healthy *Polygala myrtifolia* samples (0.2 $g.ml^{-1}$) spiked with CFBP 8416 strain suspensions ranging from $1\times10^2$ to $1\times10^6$ $CFU.mL^{-1}$. These tests allowed the determination of the limit of VNTR genotyping directly in plant extract as a function of Cq, as spiked solutions were all tested with the Harper’s qPCR test[7].

##### 1.2.3 VNTR-13 scheme genotyping

In the optimized protocol, PCRs were performed in a 20 µL-final volume containing 1X PCR buffer (no magnesium chloride) (Invitrogen), 1.5 mM of magnesium chloride (Promega), 0.2 mM of each dNTP, 0.2 µM of each primer (Promega), 1U of Platinum® Taq DNA Polymerase (Invitrogen) and 2 µL of sample DNA or 1 µL of boiled bacterial suspension. The optimal PCR conditions were 5 min at 95°C, followed by 35 cycles of 30 s at 95°C, 30 s at the annealing temperature (Table 1) and 30 s at 72°C, and ended by 10 min at 72°C.

For samples previously detected infected by *X. fastidiosa* with the Harper’s qPCR test at Cq<27, VNTRs were amplified by four in multiplex. For those with Cq values equal at 27 or lower than 32, they were amplified in simplex and PCR products multiplexed prior capillary electrophoresis analysis. Then, 2.4 µL of PCR products were mixed with 9.35 µL of Hi-Di formamide (Sigma-Aldrich) and 0.15 µL of Genescan 500 Liz internal line size standard (Applied Biosystems) for capillary electrophoresis using an ABI PRISM 3130 genetic analyzer (Applied Biosystems): injection of 16 sec at 1.2 kvolts, voltage steps 20 nk, interval of 15 sec, data delay time of 60 sec, run of 12000 sec at 15 kvolts. To test VNTR typing repeatability and reproducibility of the entire experiment, the strain CFBP 8416 was included as a control in each series of amplification and sequencing.

#### 1.3 ABC

##### 1.3.1 Scenarios

**The bottom-up approach** consisted in searching for the most probable topology for each population combination of the American ancestor and two French populations in presence or not of an unsampled (ghost) population. Altogether this lead to three independent analyses and in each analysis, 10 to 12 topologies were tested (Fig. S9 A). A total of 30 scenarios (combination of population choice and topology) were analyzed for ST6 and 36 for ST7 in this bottom-up approach.

Topologies 1, 5, 9, 10, 11 and 12 assumed that the French populations were introduced separately from two events. Topologies 2 and 6 hypothesizes that samples which belonged to the same population may have been mistakenly separated into two populations. Topologies 3, 4, 7 and 8 assumed that a first population was introduced in France and that a bridgehead invasion was responsible for the emergence of the second French populations. Topologies 5, 6, 7, 8, 10, 11 and 12 assumed that one or two populations may have been unsampled (named ghost population).

**The top-down approach** consisted in searching for the most probable population combinations for each possible topology. A total of 5 topologies were tested for each of the three French ST6 and ST7 populations (six combinations) (Fig. S9 B). The topologies differed by the number of independent introductions and divergence events. However, based on the results of the bottom-up approach, a number of scenario were considered statistically impossible and were not tested in the top-down approach.

In consequence, from the six possible population combinations all were tested for ST6, for a total of 30 scenarios (Fig. S10) and only two topologies for ST7 for a total of 12 scenarios (Fig. S11).

Class 1 topology (I) assumed three independent introductions in France from a source population. Class 2 topology (II) assumed that one population was introduced in France and is responsible for two independent bridgehead invasions. Class 3 topologies (III.I and III.II) assumed that two independent introductions occurred and one of them was responsible for a bridgehead invasion of the third population. Class 4 topology (IV) assumed that one population was first introduced and was responsible for two successive bridgehead invasions.

##### 1.3.2 Choice and validation of priors

For the DiyABC analyses, parameters were drawn in the prior distribution described in Supplementary data 8. Due to the lack of knowledge concerning the bacterium population biology (population effective sizes, date of founder event, existence and duration of a bottleneck, for example), a log uniform distribution with large ranges was chosen. Parameters for microsatellite mutation models were set by default following a stepwise mutation model[8]. Other parameters of the mutation models were set by default at “0” and all summary statistics were selected.

Prior choices were validated by performing a principal component analysis (PCA) implemented in DiyABC, in the space of summary statistics for all simulated datasets (106 per scenario) with parameter values drawn from the prior parameters. The observed dataset was projected as a supplementary individual to determine by visual check whether it fell within the variability of simulated data[8].

##### 1.3.3 Simulations and posterior probability estimation

A total of 106 simulations were performed for each competing scenario using DiyABC software[9]. Because the DiyABC lack of discriminative power among competing scenarios, abcrf were used to estimate best scenarios or the rejected one, using the R package “abcrf”[10]. Following Pudlo et al., 2016 abcrf treatments were processed on reference tables including 104 simulations per scenario[10]. In the constructed random-forests the number of trees was fixed at n=1000. For each abcrf analysis the best scenario, its prior error rates and its posterior probability were evaluated over 10 replicate runs of the same reference table. Moreover, for the bottom-up approach, an event was rejected when less than 50/1000 (<5%) of votes were attributed to it. Scripts published by Fraimout *et al*., 2017 were optimized to fit our dataset[11].

### 2. Supplementary results

#### 2.1 MLVA

##### 2.1.1 Better genotyping resolution with MLVA within *X. fastidiosa* subsp *multiplex*

*In silico* analysis of the *X. fastidiosa* subsp. *multiplex* strain M12 genome sequence led to the identification of 113 VNTR loci, with repeat units ranging from 4 bp to 44 bp in size. After further investigations on a subcollection of 46 genome sequences representing the known diversity of *X. fastidiosa* (10 *X. fastidiosa* subsp. *multiplex*, 18 *X. fastidiosa* subsp. *fastidiosa*, 18 *X. fastidiosa* subsp. *pauca*)[2]and then on a panel test of 16 strains from the present study, only three VNTR loci were retained to complete the set of 10 VNTR loci previously proposed by Lin *et al*. (2005) and Francisco *et al*. (2017) (Table 1) [3,4]. These 13 VNTR loci were present in all *X. fastidiosa* genome sequences examined, were not redundant with each other, were highly polymorphic, had a TR unit size ranging from 4 bp to 12 bp and a total array sizes from 77 to 360 bp. Among these 13 VNTR loci, all were chromosomic and six were intergenic (Table 1). Moreover, *in silico* analyses on the WGS database of bacteria from NCBI revealed that specific amplifications were exclusively obtained on all *X. fastidiosa* genome sequences and on no other genome sequence (date of analysis: 2018-08-22). The 110 other TR loci detected were not further considered, as 39 were imperfect motifs or had incomplete repetitions, 10 were redundant with previously published VNTRs, and 61 had a low allelic diversity or did not allowed to discriminate between ST6 and ST7 as examined on the 10 *X. fastidiosa* subsp. *multiplex* genome sequences.

The VNTR-13 scheme was specifically developed to type the subspecies *multiplex*. All 13 VNTR loci were successfully amplified and typed on our collection of 68 French and American *X. fastidiosa* subsp. *multiplex* strains (Supplementary data 2) and on the DNAs of 18 Spanish *X. fastidiosa* subsp. *multiplex* strains (Supplementary data 2). Interestingly, each VNTR locus was also successfully amplified on the 27 *X. fastidiosa* strains belonging to nine different sequence types of *X. fastidiosa* subsp. *fastidiosa* and *pauca* (Supplementary data 2, Fig. S5). Of particular interest, and consistent with SNP data, strains belonging to ST6 were scattered into two clusters, one grouping French and American ST6 strains, and the other grouping ST6 strains from Spain (Fig. S5). The MLVA accurately resolved the different haplotypes in our international strain collection as the haplotype accumulation curve reached a plateau with more than 95% of the haplotypes detected with nine or more markers (Fig. S6).

##### 2.2.2 Efficient genotyping of the VNTR-13 scheme on both *X. fastidiosa* subsp. *multiplex* isolated strains and DNA extracted from plants

Besides a conventional use on isolated strains, VNTR genotyping was attempted directly on large sets of DNA samples extracted from frozen infected plant material. Genotyping conditions were thus optimized by: i) multiplexing the primers by four according to their annealing temperature

(Table 1); ii) selecting a highly efficient DNA polymerase, meaning the least affected by PCR inhibitors. Platinum™ Taq DNA Polymerase (ThermoFisher) was selected as it allowed amplification of some VNTR loci (ASSR-9, ASSR-16, GSSR-4) in bacterial suspensions at ten times lower concentrations than GoTaq® G2 Flexi DNA Polymerase (i.e. concentrations as low as $10^3$ to $10^4$ CFU.mL$^{-1}$); and iii) setting sample DNA aliquot volume to 2 µL per 20 µL-total volume reaction to maximize target presence and amplification, as no inhibitory effect was noticed (data not shown).

The MLVA proved to have an excellent ability to genotype *X. fastidiosa* from isolated strains as well as from DNA extracted from plant sample. In fact, we had at our disposal 13 pairs of isolated strains and DNAs extracted from the same original plant samples for a total of five different plant species (*Acacia dealbata*, *Euryops chrysanthemoides*, *Medicago sativa*, *Polygala myrtifolia*, and *Prunus avium*) (Supplementary data 3). An identical VNTR profile was obtained for each member of these 13 pairs, indicating that VNTR amplification from DNA extracted from plant material was efficient and was not affected by DNA extraction efficiency and/or quality. From this point, each of these 13 pairs was considered as one sample.

##### 2.2.3 Population structure of the French *X. fastidiosa* subsp. *multiplex* strains

Because DAPC and STRUCTURE analyses led to almost identical groupings (Fig. 3, Fig. S8), the multivariate analysis of population subdivision method of the DAPC was used, as it does not rely on an underlying population genetics model[12], in contrast to the model-based clustering algorithm of STRUCTURE. As expected, ST6 and ST7 strains from plant samples were always separated in different clusters. We tested increasing numbers of clusters from k=2 upwards, and for k>6, F-statistics ($R_{ST}$ and $F_{ST}$) did not validate the groupings. For k=4, cluster 1 was mainly composed of ST6 samples from Corsica (135 Corsica and two PACA, in turquoise Fig. 3); cluster 2 grouped the rest of the 47 ST6 samples (19 Corsica and 28 PACA, in red Fig. 3); cluster 3 was mainly composed of the ST7 samples from Corsica (110 Corsica and eight PACA, in blue Fig. 3); and cluster 4 grouped the remaining 94 ST7 samples mainly sampled from PACA (19 Corsica and 75 PACA, in yellow Fig. 3). In the 20 independent DAPC runs, all 396 samples, except 16, were consistently assigned to a single DAPC cluster (Supplementary data 3). For these 16 remaining samples, the 11 ST6 samples, all from PACA, were most often (i.e. in 70% of the runs) assigned to DAPC cluster 2 (mix geographical origin), and less frequently to DAPC cluster 1 (i.e. in 30% of the runs). The five ST7 samples from Corsica and PACA were most often (i.e. in 80 % of the runs) assigned to DAPC cluster 4, and less frequently to DAPC cluster 3 (i.e. in 20 % of the runs). All samples grouped in clonal complexes by MST analysis clustered together at k=4 by DAPC analysis. For lower and higher values of k (two to six) each ST6 and ST7 group of samples were gradually divided from one to three clusters. At k=6, two of the three

ST6 clusters grouped both samples isolated in Corsica and PACA regions, without any obvious clustering in relation to host or year of isolation. Moreover, comparing these clusters and the MST, the main ST6 clonal complex containing 95 samples was divided into two clusters. Regarding ST7, the three clusters grouped samples isolated from both regions. MST algorithm and DAPC and STRUCTURE provided coherent clustering for k=4 and this clustering made biological sense (clustering mainly coherent with geographical origin of the strain or potential dissemination of haplotypes), so it was used as a first basis for further analyses.

**Fig. S1. Genome clustering by FastSTRUCTURE software for K=5.** Color bars represent groups identified by Bayesian clustering. ST numbers are indicated above strain codes. Color names represents their country of origin of strains: black for France, blue for USA, green for Brazil, purple for Italy and red for Spain.

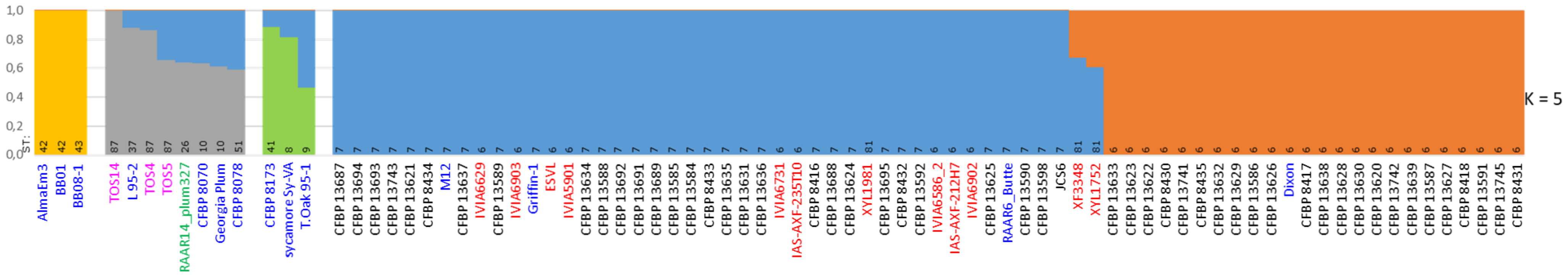

**Fig. S2. Heatmap representing the number of SNPs that came from mutations and differed between the 82 genome sequences**

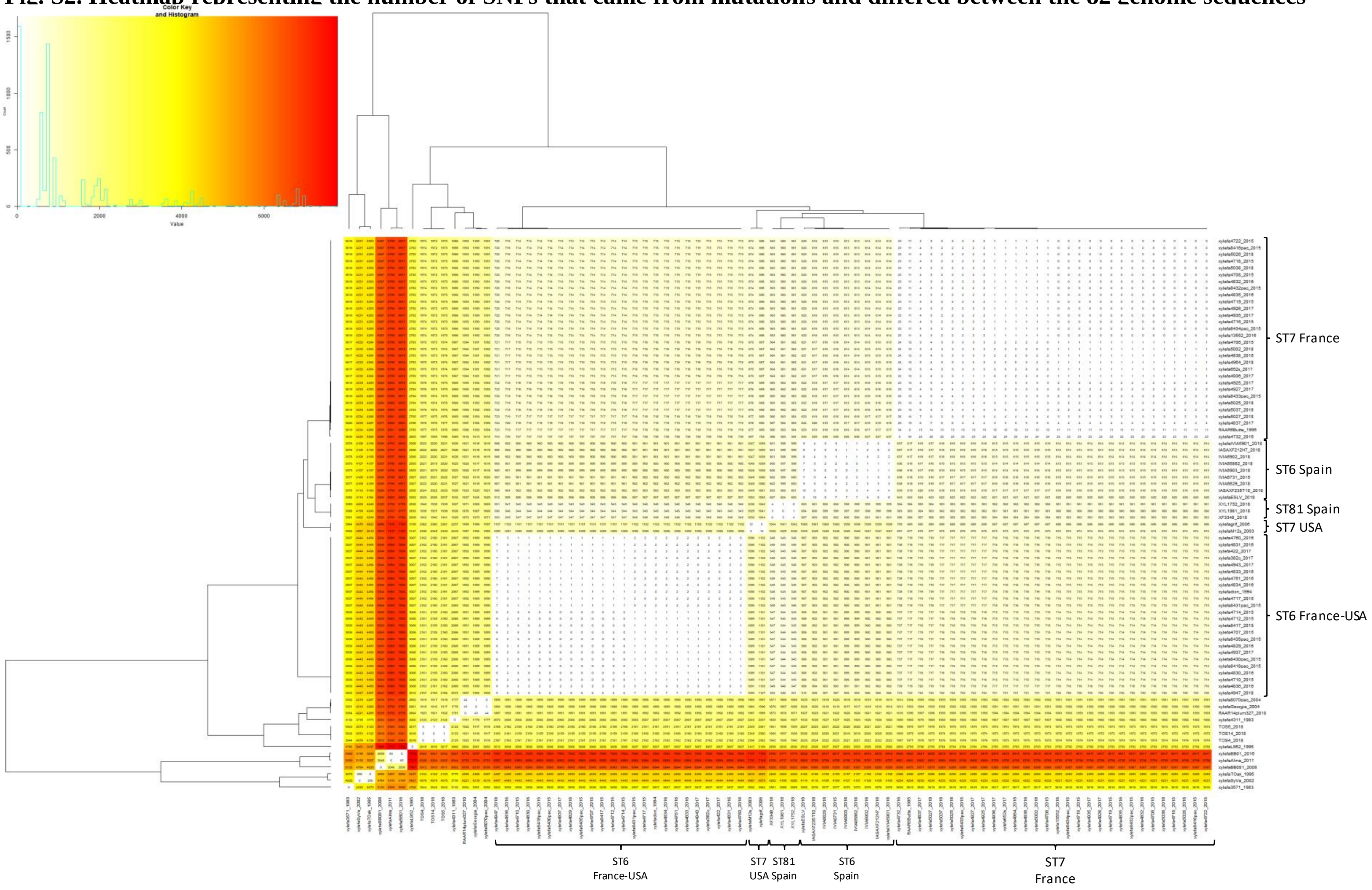

**Fig. S3. Local temporal signal detection.** Linear regression between the age of the samples and their root-to-tip genetic distances.

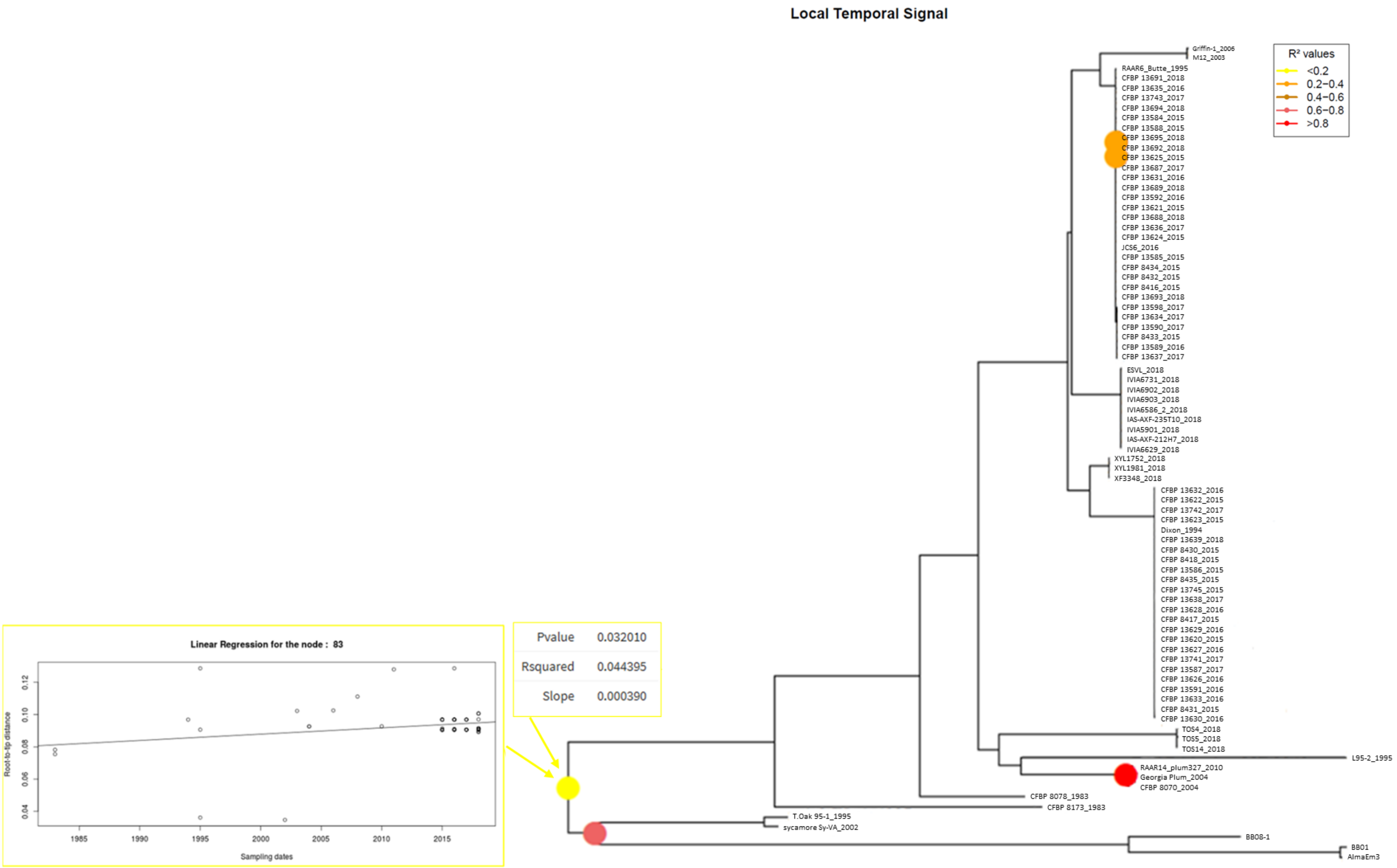

**Fig. S4. Date randomization tests.** Comparison of A) the mean rate, B) the ucldMean, C) the tree height estimated using BEAST 2.6.1. from the original dataset (red) and 40 date-randomized datasets using the TIPDATINGBEAST R package.

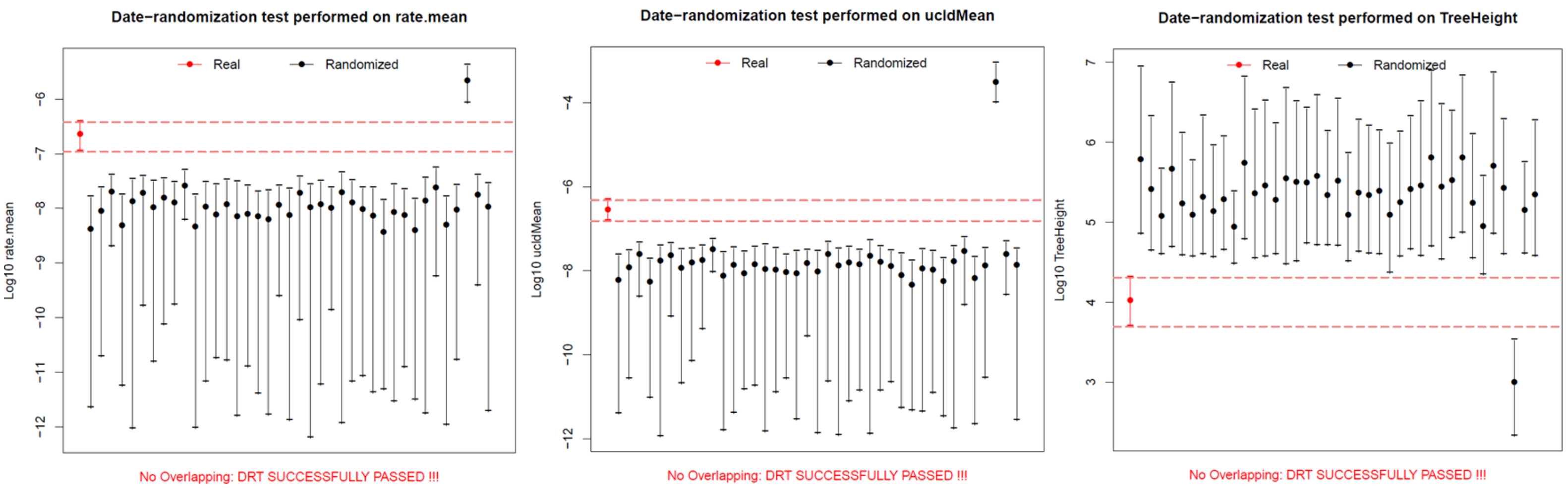

**Fig. S5. Minimum Spanning Trees of the 113 *X. fastidiosa* strains isolated in Costa Rica, Ecuador, France, the Netherlands, Spain and the USA typed using the VNTR-13 scheme.** Colors refer to A) the sequence type of the strains, B) the country of origin of the strains. For details on strains refer to Supplementary data 2 and 3.

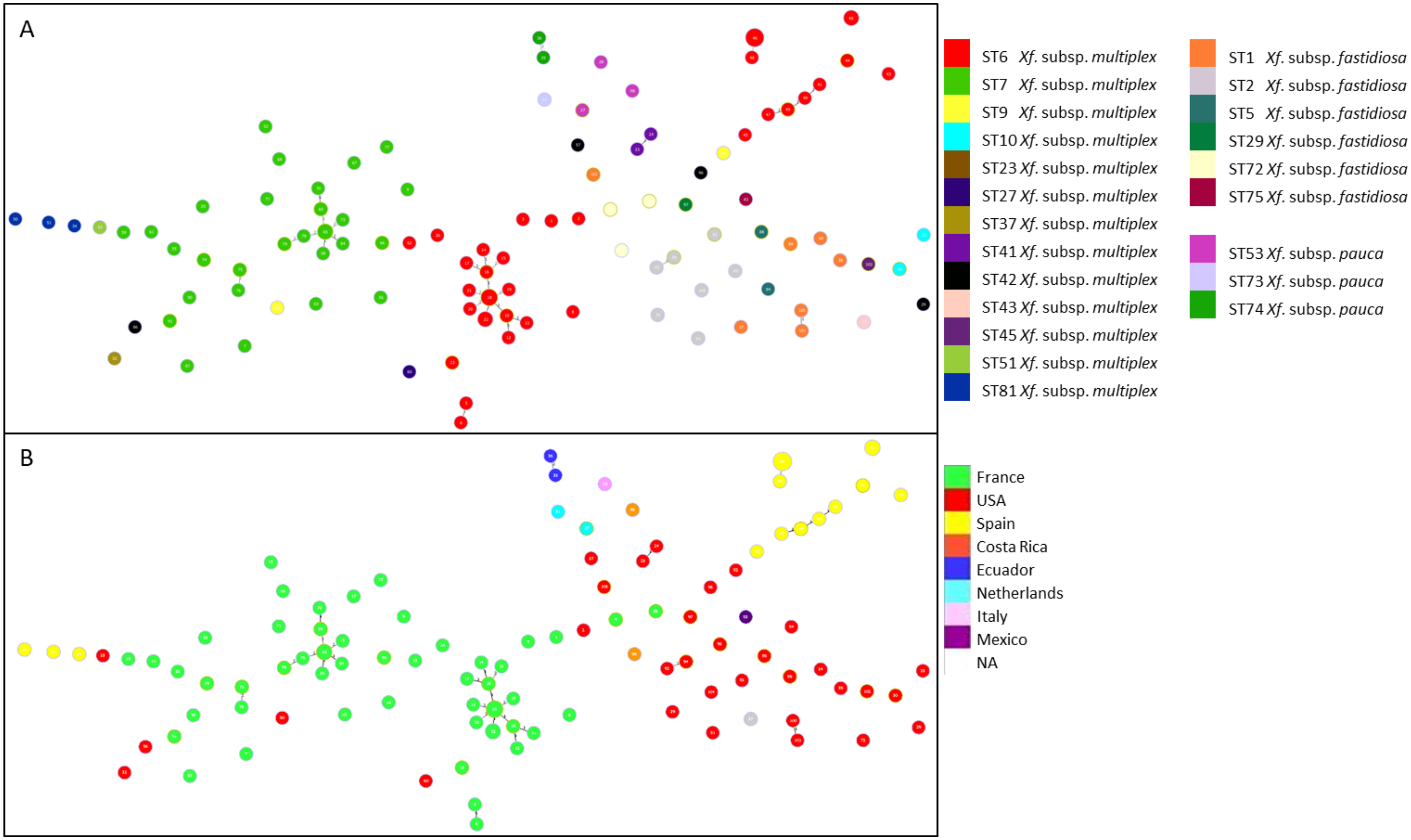

**Fig. S6. Genotype accumulation curve of X. fastidiosa over 13 loci. A) for the 113 strains of the study, B) on the 396 French samples.**

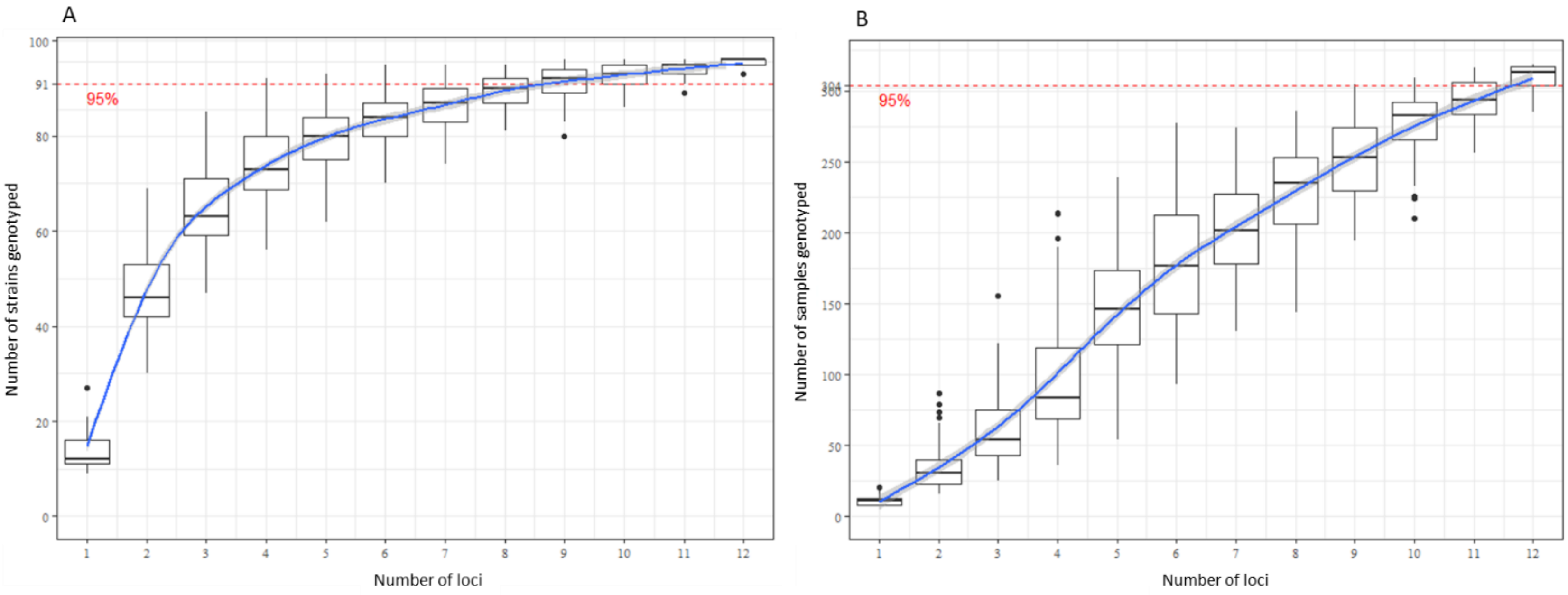

**Fig. S7. Allele frequencies at each VNTR locus per sequence type for the 396 French X. fastidiosa infected samples.** At each locus, colors represent the set of haplotypes.

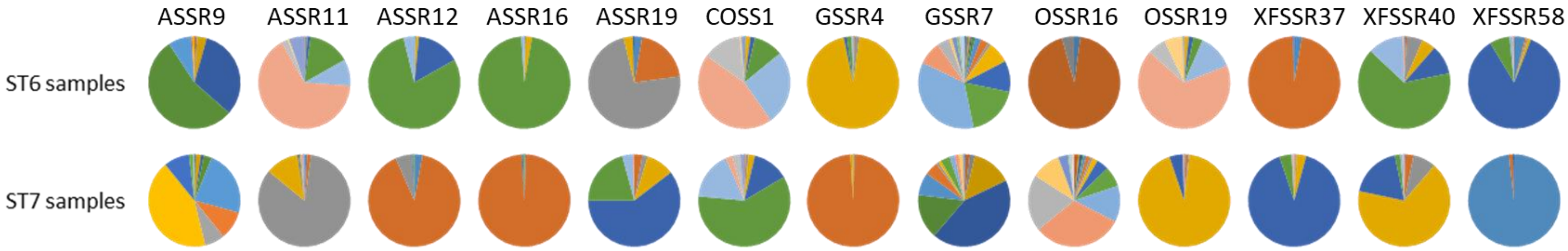


**Fig. S8. Sample clustering by STRUCTURE software. Colors represent groups identified by Bayesian clustering.** Cluster 1 (turquoise), cluster 2 (red), cluster 5 (orange) grouped ST6 samples and cluster 3 (blue), cluster 4 (yellow), cluster 6 (purple) grouped ST7 samples.

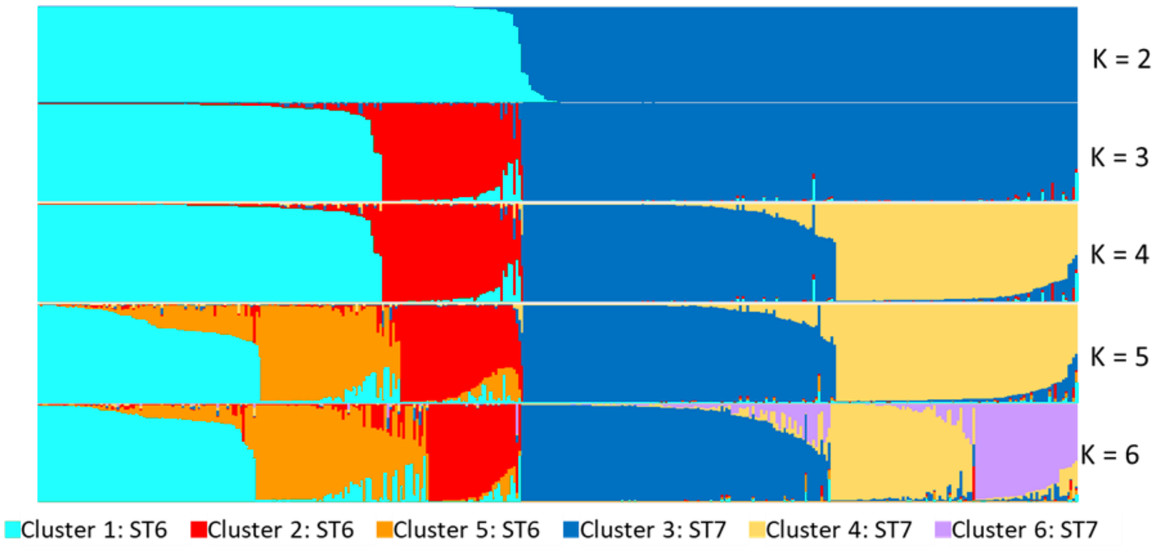

**Fig. S9. The scenarios compared using ABC used independently for ST6 and ST7 samples.** A) The 12 scenarios compared by the bottom-up approach. All scenarios considered that the French populations (Pop1 and Pop2) derived from an initial USA population. Ghost populations, *i.e.* populations that would have existed but were not sampled, were used in scenario 5-8 and 10-12. The time on the Y axis refers to the divergence date between populations. B) The five topologies analyzed by the top-down approach. All scenarios considered that the French populations (Pop1, Pop2 and Pop3) derived from an initial USA population. The time on the Y axis refers to the divergence date between populations.

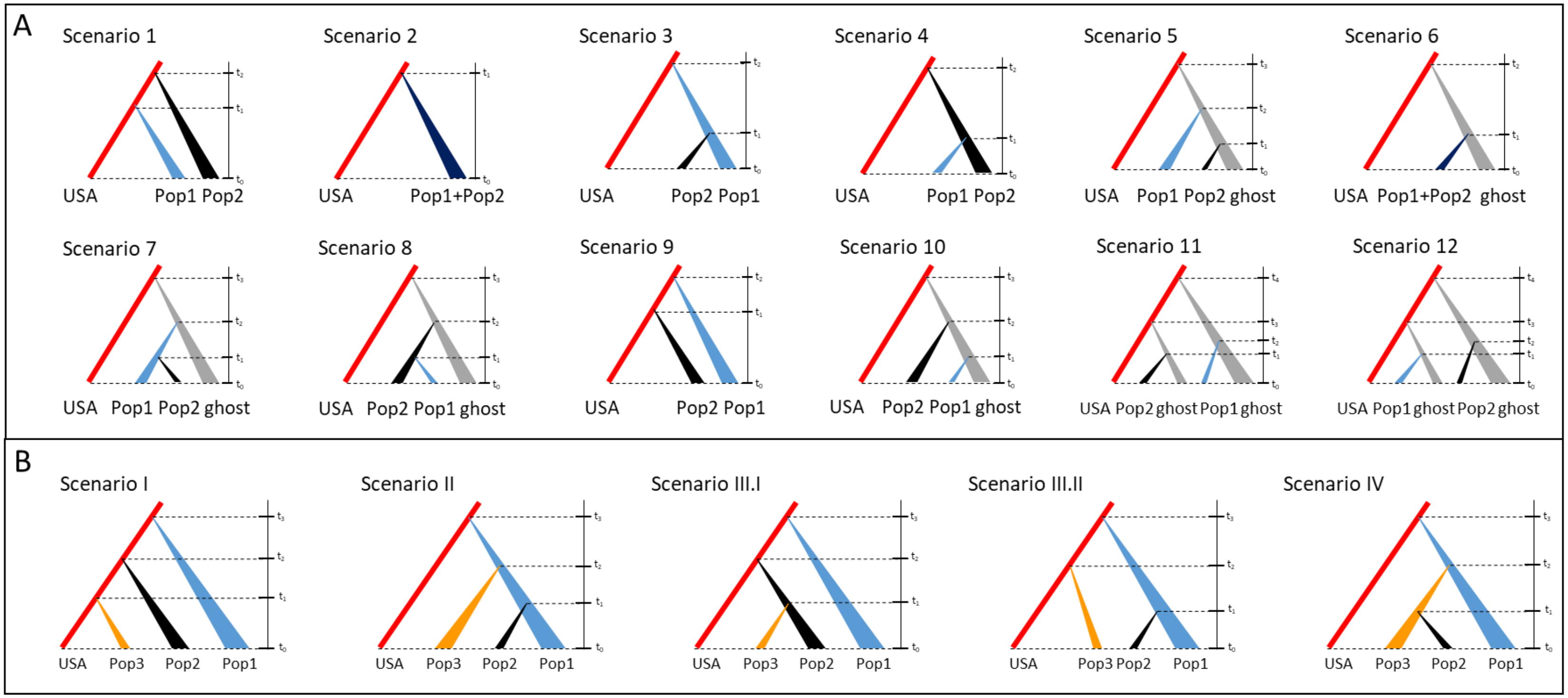

**Fig. S10. List and visual representation of the 30 scenarios analyzed by ABC for ST6 samples.** All scenarios considered that the French populations (C1P1, C2, and P2) derived from an initial USA population. The time on the Y axis refers to the divergence date between populations. Scenario II.7 (boxed in red) is the one defined by Abcrf analysis as the most probable.

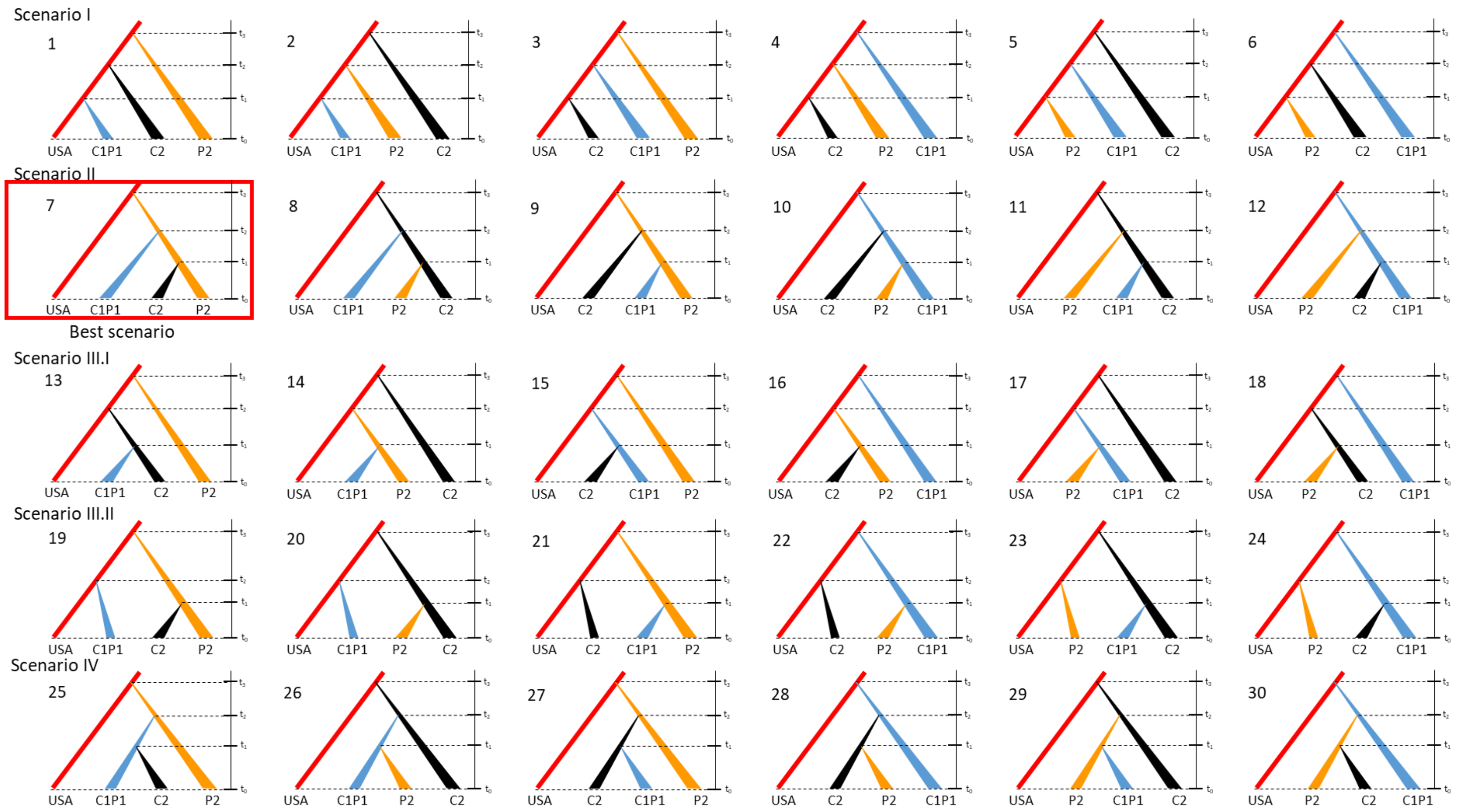

**Fig. S11. List and visual representation of the 12 scenarios analyzed by ABC for ST7 samples.** All scenarios considered that the French populations (C1P1, C2, and P2) derived from an initial USA population. The time on the Y axis refers to the divergence date between populations. Scenario II.7 (boxed in red) is the one defined by Abcrf analysis as the most probable.

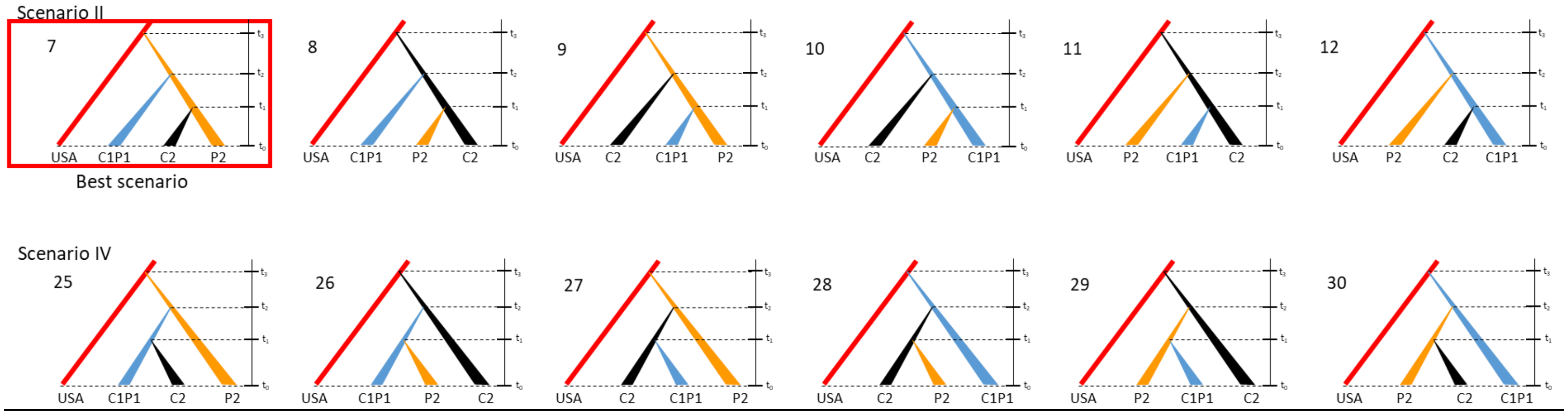

**References Supplementary Materials**


1. Mayjonade, B. *et al.* Extraction of high-molecular-weight genomic DNA for long-read sequencing of single molecules. *BioTechniques* **61**, 203–205 (2016).

2. Denancé, N., Briand, M., Gaborieau, R., Gaillard, S. & Jacques, M.-A. Identification of genetic relationships and subspecies signatures in Xylella fastidiosa. *BMC Genomics* **20**, 1–21 (2019).

3. Lin, H. *et al.* Multilocus simple sequence repeat markers for differentiating strains and evaluating genetic diversity of Xylella fastidiosa. *Applied and environmental microbiology* **71**, 4888–4892 (2005).

4. Francisco, C. S., Ceresini, P. C., Almeida, R. P. P. & Coletta-Filho, H. D. Spatial Genetic Structure of Coffee-Associated Xylella fastidiosa Populations Indicates that Cross Infection Does Not Occur with Sympatric Citrus Orchards. *Phytopathology* **107**, 395–402 (2017).

5. Koressaar, T. & Remm, M. Enhancements and modifications of primer design program Primer3. *Bioinformatics* **23**, 1289–1291 (2007).

6. Engels, W. R. Contributing software to the internet: the amplify program. *Trends in Biochemical Sciences* **18**, 448–450 (1993).

7. Harper, S. J., Ward, L. I. & Clover, G. R. G. Development of LAMP and real-time PCR methods for the rapid detection of Xylella fastidiosa for quarantine and field applications. *Phytopathology* **100**, 1282–1288 (2010).

8. Cornuet, J.-M., Ravigné, V. & Estoup, A. Inference on population history and model checking using DNA sequence and microsatellite data with the software DIYABC (v1.0). *BMC Bioinformatics* **11**, 401 (2010).

9. Cornuet, J.-M. *et al.* Inferring population history with DIY ABC: a user-friendly approach to approximate Bayesian computation. *Bioinformatics* **24**, 2713–2719 (2008).

10. Pudlo, P. *et al.* Reliable ABC model choice via random forests. *Bioinformatics* **32**, 859–866 (2016).

11. Fraimout, A. *et al.* Deciphering the routes of invasion of Drosophila suzukii by means of ABC random forest. *Mol Biol Evol* msx050 (2017) doi:10.1093/molbev/msx050.

12. Jombart, T., Devillard, S. & Balloux, F. Discriminant analysis of principal components: a new method for the analysis of genetically structured populations. *BMC Genetics* **11**, 1–15 (2010).

**References Supplementary Tables**

Bhattacharyya, A. *et al.* Whole-genome comparative analysis of three phytopathogenic Xylella fastidiosa strains. *Proc. Natl. Acad. Sci. U.S.A.* **99**, 12403–12408 (2002).

Chen, J. *et al.* Whole Genome Sequences of Two Xylella fastidiosa Strains (M12 and M23) Causing Almond Leaf Scorch Disease in California. *J Bacteriol* **192**, 4534 (2010).

Van Horn, C., Chang, C.-J. & Chen, J. De Novo Whole-Genome Sequence of Xylella fastidiosa subsp. multiplex Strain BB01 Isolated from a Blueberry in Georgia, USA. *Genome Announc* **5**, e01598-16 (2017).

Giampetruzzi, A. *et al.* Draft Genome Resources of Two Strains (“ESVL” and “IVIA5901”) of Xylella fastidiosa Associated with Almond Leaf Scorch Disease in Alicante, Spain. *Phytopathology* **109**, 219–221 (2019).

Chen, J., Huang, H., Chang, C.-J. & Stenger, D. C. Draft Genome Sequence of Xylella fastidiosa subsp. multiplex Strain Griffin-1 from Quercus rubra in Georgia. *Genome Announc* **1**, e00756-13 (2013).

Giampetruzzi, A. *et al.* Draft Genome Sequence Resources of Three Strains (TOS4, TOS5, and TOS14) of Xylella fastidiosa Infecting Different Host Plants in the Newly Discovered Outbreak in Tuscany, Italy. *Phytopathology* **109**, 1516–1518 (2019).
